# Engineering Human CNS Morphogenesis: Controlled Induction of Singular Neural Rosette Emergence

**DOI:** 10.1101/229328

**Authors:** Gavin T. Knight, Brady F. Lundin, Nisha Iyer, Lydia M.T. Ashton, William A. Sethares, Rebecca L. Willett, Randolph S. Ashton

**Affiliations:** Department of Biomedical Engineering, University of Wisconsin-Madison, Madison, WI 53706, USA; Wisconsin Institute for Discovery, University of Wisconsin-Madison, Madison, WI 53715, USA; Department of Consumer Science, University of Wisconsin-Madison, Madison, WI 53706, USA; Department of Electrical and Computer Engineering, University of Wisconsin-Madison, Madison, WI 53706, USA

**Keywords:** neural organoids, neuroepithelium, micropatterning, click chemistry

## Abstract

Human pluripotent stem cell (hPSC)-derived neural organoids have revolutionized in vitro modelling of human neurological disorders. Cell-intrinsic morphogenesis processes displayed within these tissues could serve as the basis for ex vivo manufacture of brain and spinal cord tissues with biomimetic anatomy and physiology. However, we must first understand how to control their emergent properties starting at the genesis of neural organoid formation, i.e. emergence of polarized neuroepithelium. In vivo, all CNS tissues develop from a singular neuroepithelial tube. Yet, current protocols yield organoids with multiple neuroepithelial rings, a.k.a. neural rosettes, each acting as independent centers of morphogenesis and thereby impeding coordinate tissue development. We discovered that the morphology of hPSC-derived neural tissues is a critical biophysical parameter for inducing singular neural rosette emergence. Tissue morphology screens conducted using micropatterned array substrates and an automated image analysis determined that circular morphologies of 200-250 and 150μm diameter are optimal for inducing singular neural rosette emergence within 80-85% forebrain and 73.5% spinal tissues, respectively. The discrepancy in optimal circular morphology for Pax6^+^/N-cadherin^+^ neuroepithelial forebrain versus spinal tissues was due to previously unknown differences in ROCK-mediated cell contractility. The singular neuroepithelium induced within geometrically confined tissues persisted as the tissues morphed from a 2-D monolayer to multilayered 3-D hemispherical aggregate. Upon confinement release using clickable micropatterned substrates, the tissue displayed radial outgrowth with maintenance of a singular neuroepithelium and peripheral neuronal differentiation. Thus, we have quantitatively defined a pertinent biophysical parameter for effectively inducing a singular neuroepithelium emergence within morphing hPSC-derived neural tissues.

**Significance Statement:** Human pluripotent stem cell (hPSC)-derived neural organoids display emergent properties that, if harnessed, could serve as the basis for ex vivo manufacture of brain and spinal cord tissues with biomimetic macroscale anatomy and physiology. Their chaotic terminal structure arises from uncontrolled morphogenesis at their genesis, resulting in spontaneous induction of multiple neuroepithelial morphogenesis centers,a.k.a. neural rosettes. Here, we determined that neural tissue morphology is a pertinent biophysical parameter for controlling subsequent morphogenesis, and identified discrete circular tissue morphologies as optimal and effective at inducing singular neural rosette emergence within forebrain and spinal neural tissues. Thus, we developed an approach to reproducibly control the initial stage of hPSC-derived neural tissue morphogenesis enabling their manufacture with a biomimetic nascent CNS anatomy.

## Introduction

The derivation of organoids from human pluripotent stem cells (hPSCs) has redefined the possibilities of in vitro tissue engineering. Human PSCs have traditionally been characterized by their ability to spontaneously recapitulate facets of developmental tissue morphogenesis upon ectopic implantation in vivo, i.e. the teratoma assay (1). However, more recent discovery of their ability to also self-organize emergence of organotypic tissues in vitro, e.g. cortical (2), retinal (3), cerebral (4) and intestinal organoids (5), has spurred the creation of novel developmental and disease models with unprecedented biomimicry of microscale cytoarchitectures and cell phenotype diversity (6, 7). The innate emergent properties of hPSCs could serve as the basis for advanced biomanufacture of functional tissue and organ models or even transplants (5, 8). Yet, it remains a challenge to engineer reproducible hPSC morphogenesis in vitro and thereby enable controlled emergence of organotypic tissues with standardized cytoarchitecture (9, 10).

Morphogenesis of the human central nervous system (CNS) as well as hPSC-derived neural organoid commences with neurulation of columnar neuroepithelial cells (NECs). In this dynamic process, NECs polarize adherens, e.g. N-cadherin, and tight junction proteins towards an apical lumen while depositing extracellular matrix (ECM) proteins at their basal surface. In vivo, neurulation results in emergence of a singular neuroepithelial tube that spans the entire rostrocaudal (R/C) axis of the embryo’s dorsal plane and serves as the primordium of all brain, retinal, and spinal cord tissues. This singular polarized neuroepithelium serves as a critical morphogenesis center for organizing establishment of lamellar tissue cytoarchitectures during subsequent stages of CNS development. For example, it localizes waves of mitotic NEC proliferation at the tube’s apical surface while daughter cells migrate radially towards the basal surface to complete differentiation, functional maturation, and expand the CNS parenchyma. Disruption of the neuroepithelial tube’s N-cadherin polarized cytoarchitecture eliminates the sole neurogenic source of embryonic CNS development (11). Alternatively, the presence of multiple neural tubes during development causes congenital abnormalities such as dipolmyelia and diastematomyelia (12, 13) or anencephaly as observed in diprosopic parapagus twins (14). Therefore, formation of singular neuroepithelial tissues within hPSC-derived organotypic tissues is an important cytoarchitectural feature for generating biomimetic CNS tissue models (15).

In current 2- and 3-D hPSC-derived cultures, neurulation occurs spontaneously yielding uncontrolled emergence of numerous polarized neuroepithelial tissues, a.k.a. neural rosettes (4, 16). Integration of engineering techniques such as stirred-tank bioreactor culture during 3-D neural organoid derivation can induce the formation of larger and more contiguous neuroepithelial tissues (4, 17). Likewise, derivation of neural organoids around filamentous biomaterial scaffolds was shown to induce elongated neuroepithelial tissues within the 3-D organoid (10). Yet, the persistent presence of multiple neural rosettes of indiscriminate shapes and sizes within a single organotypic tissue inevitably confounds subsequent morphogenesis events, potentially limiting tissue maturation and producing significant variability in the resultant cytoarchitecture. Interestingly, singular neural rosette emergence was routinely observed upon derivation of 3-D neuroepithlelial cysts, which have considerably smaller dimensions than hPSC-derived organoids and are generated from single cell mouse embryonic stem cell (mESC) suspensions (18, 19). Manual isolation of singular neural rosettes from culture remains the only reliable in vitro method for generating such critical biomimetic cytoarchitecture within hPSC-derived organotypic CNS tissues (15, 20).

Geometric confinement of hPSC tissues on 2-D micropatterned substrates was recently shown to induce self-organized embryonic patterning, i.e. gastrulation, in a morphology dependent manner (21, 22). Based on this and previously discussed mESC-derived neuroepithlelial cysts studies, we investigated whether engineering the morphology of neurally differentiating hPSC tissues could similarly regulate their morphogenesis specifically neural rosette emergence. In both standard well-plate and 2-D micropatterned culture, neural rosette formation was observed to be a function of local cell density and acquisition of a Pax6^+^/N-cadherin^+^ neuroectodermal fate. Using a custom image analysis algorithm and machine learning classifier, screens of hPSC-derived neuroepithelial tissues of various morphologies revealed an indirect correlation between the tissue’s surface area and the propensity for singular neural rosette emergence. Circular micropatterns of 200-250μm diameter (0.031-0.049mm^2^) were observed to most effectively induce singular neural rosette emergence within forebrain neuroepithelial tissues reaching levels of 80-85% efficiency when initially seeded as hPSCs. These studies were further extended to develop an analogous protocol for neuroepithelial tissues of a ventral spinal cord regional phenotype. This revealed that spinal neuroepithelial tissues, unlike their forebrain counterparts, preferentially formed singular neural rosettes (73.5% efficiency) on circular micropatterns of a smaller 150μm diameter (0.018mm^2^), indicating novel biophysical differences between NECs of varying regional phenotypes. During the rosette emergence period, the proliferative neural tissues morphed from a 2-D monolayer to a 3-D, multi-layered, disk morphology of the prescribed micropatterned diameter. Upon release from micropattern confinement, the tissue slices expanded radially while maintaining a singular neuroepithelial rosette as the morphogenesis center, depositing a basal lamina, and displaying neuronal differentiation at the tissues’ periphery. Thus, we have developed a culture platform for engineering controlled neural rosette emergence within hPSC-derived organotypic CNS tissues. Reproducible induction of this nascent cytoarchitecture is a pre-requisite for advanced biomanufacture of human CNS organoids with consistent, biomimetic anatomy (9).

## Results

### Characterization of neural rosette emergence in well plate culture

Forebrain NECs were derived from hPSCs using our previously published E6 protocol (23). In six days and without the use of SMAD inhibitors (24), the E6 protocol generates near homogenous monolayers of Pax6^+^/Sox2^+^/N-cadherin^+^ NECs that undergo spontaneous neurulation to produce cultures densely populated with neural rosettes of varying shapes and sizes (**Fig. 1A**). To better characterize neural rosette emergence, we conducted a time course analysis using Pax6 and N-cadherin immunostaining (**Fig. 1B)**. In agreement with previous studies (23), the onset of Pax6 expression, a human neuroectodermal fate determinant (25), began between days 2 (0%) and 3 (44±15%) of the E6 protocol (**Fig. 1C-D**). By day 3 and concurrent with increasing Pax6 expression, the presence of polarized N-cadherin^+^ foci were observed. By day 5, Pax6 expression had reached 81±5% and N-cadherin^+^ polarization foci had both increased in prevalence and morphed into coherent rings surrounded by contiguous alignment of polarized NECs, which is characteristic of neural rosettes. Thus, neural rosette emergence occurs over five days after commencing E6 culture in 6-well plates with correlated and progressive increases in Pax6 expression and N-cadherin polarization.

**Fig. 1.**
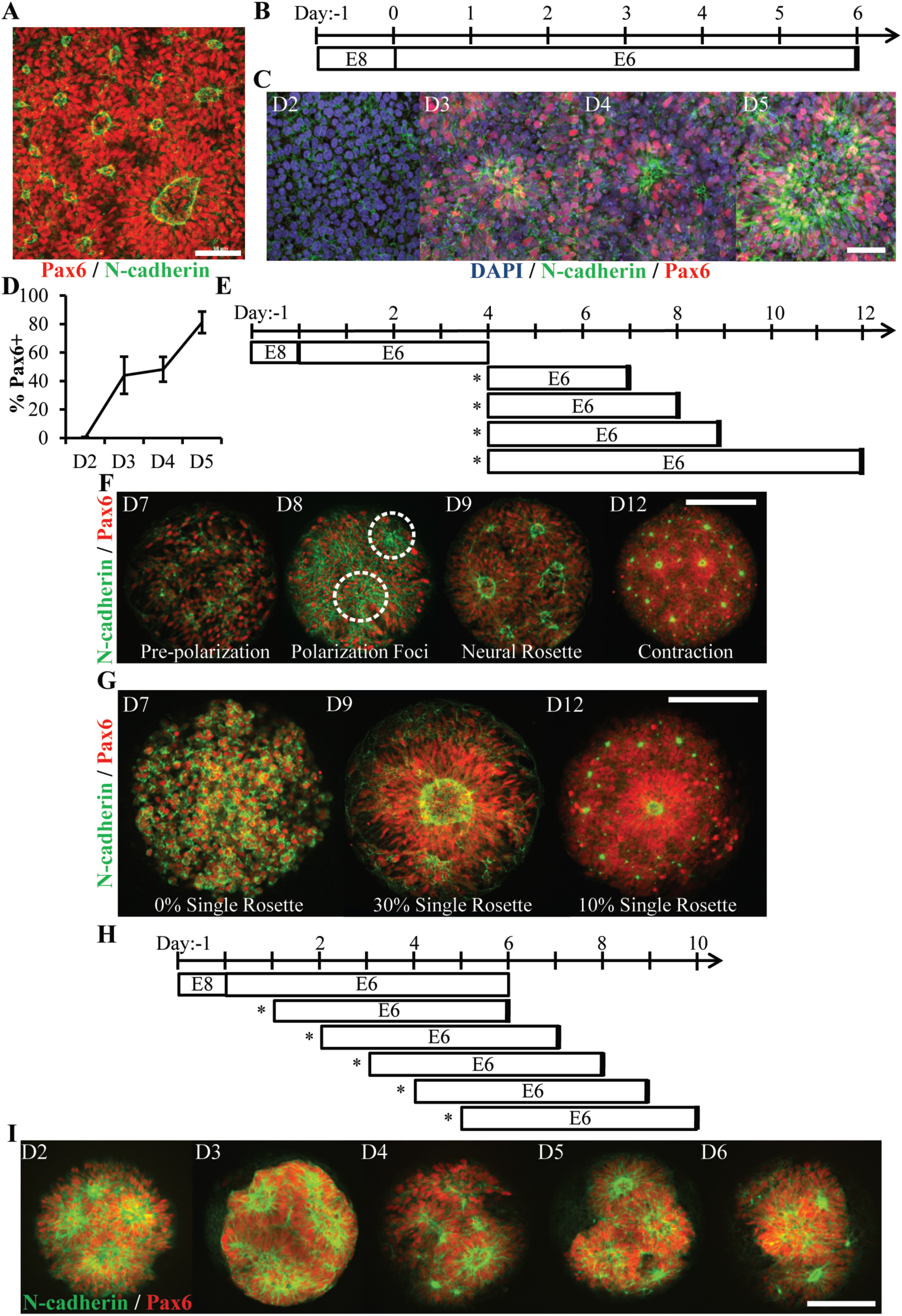
Characterization of Neural Rosette Emergence. (A) Image of N-cadherin polarized neural rosettes structures within neuroepithelial tissues derived from hPSCs using the E6 method in 6-well TCPS plates. (B) Schematic of NEC derivation via the E6 method and (C) representative immunostained images of neural induction (Pax6) and polarization (N-cadherin) time course. (D) Quantification of Pax6 expression during NECs derivation (n=6 fields of view for one well per time point). Error bars represent standard deviation. (E) Schematic of neural rosette emergence within micropatterned tissues derived from D4 NECs; sub-culture/seeding onto micropatterned substrates indicated by (*). (F) Representative images depicting stages of rosette emergence and (G) rate of singularity within neuroepithelial tissues on select days. Dotted lines highlight polarization foci. (H) Schematic of experiment testing the effect of E6 subculture time point on neural rosette emergence within micropatterned neuroepithelial tissues, and (I) corresponding images of immunostained tissues 5 days after culture on micropatterned substrates. Scale bars are (A, C) 50μm and (F, G, and I) 200μm.

### Characterization of neural rosette emergence time course within micropatterned tissues

Neural rosettes are considered an analog to 3-D neuroepithelial tube formation. Yet unlike emergence of a singular neural tube during normal development, myriad neural rosettes form spontaneously and randomly upon NEC derivation in vitro (**Fig. 1A**). We hypothesized that the prevalence of neural rosettes could be regulated by controlling tissue size and geometry. This was corroborated by our previous observation that ∼0-4 neural rosettes routinely form within micropatterned, 300μm diameter circular NEC tissues (26). Still, it remained uncertain whether engineering NEC tissue morphology could instruct reproducible emergence of a singular human neural rosette analogous to neural tube development.

Based on the temporal dynamics of rosette formation observed in well plate culture (**Fig. 1B-D**), we first characterized the timeline of neural rosette emergence within micropatterned neural tissues. NECs were harvested on day 4 of the E6 protocol (D4 NECs), which corresponded with increasing Pax6 expression and rosette formation capacity (**Fig. 1E**). Next, they were seeded at 75,000 cells/cm^2^ onto micropatterned, poly(ethylene glycol methyl ether)-grafted substrates presenting arrays of 300μm diameter circular regions (0.071mm^2^) coated with Matrigel (**Fig. S1A**) (27, 28). Time course analysis over the next 8 days using Pax6 and N-cadherin immunostaining revealed several distinct stages of rosette emergence that we classified as prepolarization (∼day 7), polarization foci (∼day 8), neural rosette (∼day 9), and contraction (∼day 13) (**Fig. 1F**). Interestingly, despite being D4 NECs upon seeding onto micropatterned substrates, ∼5 days of additional E6 culture was still required for neural rosette emergence similar to well plate culture (**Fig. 1C**). The 5 day time point also correlated with the highest occurrence (30%, n=100 tissues) of singular rosette emergence within micropatterned NEC tissues (**Fig. 1G**). Additionally, the NECs appeared to proliferate within micropatterned tissues after seeding as indicated by increased Pax6^+^ cells density (e.g. day 7 vs 9, **Fig. 1F**).

Persistence of the approximate 5-day time course for neural rosette emergence in E6 media, whether seeding hESCs at 150,000 cells/cm^2^ in well plates or D4 NECs at 75,000 cells/cm^2^ on micropatterned substrates, indicates that temporal aspects of rosette emergence may rely on a conserved aspect of E6 culture. As a final test of the 5-day paradigm, we investigated whether seeding Pax6^+^ cells on micropatterned substrates was requisite for adherence to the 5 day rosette emergence time course. Neurally differentiating cultures were subcultured at day 1, 2, 3, 4, and 5 of the E6 protocol, i.e. Pax6 expression being absent in D1-2 and present in D3-5 cultures. The cells were re-seeded onto 300μm diameter (0.071mm^2^) circular micropattern arrays. Cultures were maintained in E6 media for 5 days prior to fixation and immunostaining (**Fig. 1H**). In all conditions, the cells proliferated and produced neuroepithelial tissues with at least a 2-fold increase in cell density, >80% Pax6 expression, and consistent neural rosette formation (**Fig. 1I** and **Fig. S1B-C**). Of note, all tested cell phenotypes were observed to initially adhere as monolayers in well plates or on micropatterned substrates even when the seeding density was increased by several folds (data not shown). These results indicate that a 5-day paradigm for neural rosette emergence in E6 media holds for cultures seeded at all stages of hESC neural induction and appears to be a function of Pax6 expression and local cell density. From here onward, a 5-day micropatterned culture period was used to assess neural rosette emergence.

### Neural rosette emergence is regulated by tissue morphology

The previous experiments demonstrated that switching from a 9.6cm^2^ (a well in a 6-well plate) to a 0.071mm^2^ (300μm diameter) circular neuroepithelial tissue can substantially reduce the number of emerging neural rosettes. However, formation of neuroepithelial tissues with a biomimetic, singular rosette was not highly reproducible (30%, **Fig. 1G**). Biophysical studies have inextricably linked developmental morphogenesis events such as neural tube formation with tissue biomechanics (29, 30). Specifically, studies on micropatterned cultures have observed that multicellular tissues contract as a unit generating gradients of mechanical stresses with maximal cytoskeletal forces arising at their periphery and in spatial distributions dependent upon tissue morphology (31). Therefore, we hypothesized that the reproducibility of singular rosette emergence could be further improved by varying neuroepithelial tissue morphology, i.e. size and geometry.

This hypothesis was tested by assessing singular rosette induction efficiency within micropatterned neuroepithelial tissues of circular, triangle, square, and oblong morphologies of varying sizes (**Fig. 2A**). Culture substrates were designed to normalize the surface area of all tissue morphologies of a given size scale to 200 (0.031mm^2^), 300 (0.071mm^2^), or 400μm (0.126mm^2^) diameter circular micropatterns. In this manner, the cell seeding density would be constant amongst tissues of differing morphologies within the same size scale. Day 4 NECs were seeded onto 0.031, 0.071, and 0.126mm^2^ micropattern arrays at 33,300, 75,000, and 133,000 cells/cm^2^, respectively, to correspond with changes in micropattern cell-adhesive area across each size scale. After 5 days in E6 media, the cultures were fixed, immunostained for Pax6 and N-cadherin, and analyzed manually to quantify the number of N-cadherin^+^ polarization foci and rosettes (**Fig. 2B**). Each tissue was categorized as follows: the presence of 0 or 1 polarization foci were ‘0 Rosette’; 1 rosette and 0 polarization foci were ‘1 Rosette’; ≥ 1 rosette with ≥ 1 additional polarization foci were ‘+1 Rosette”. Although all micropatterned tissue morphologies exhibited the capacity to induce ‘1 Rosette’ neuroepithelial tissues, 0.031mm^2^ circular (56%) and square (57%) morphologies were the most efficient. Yet, manual analysis of each image limited our ability to screen larger sample sizes per tissue morphology.

**Fig. 2.**
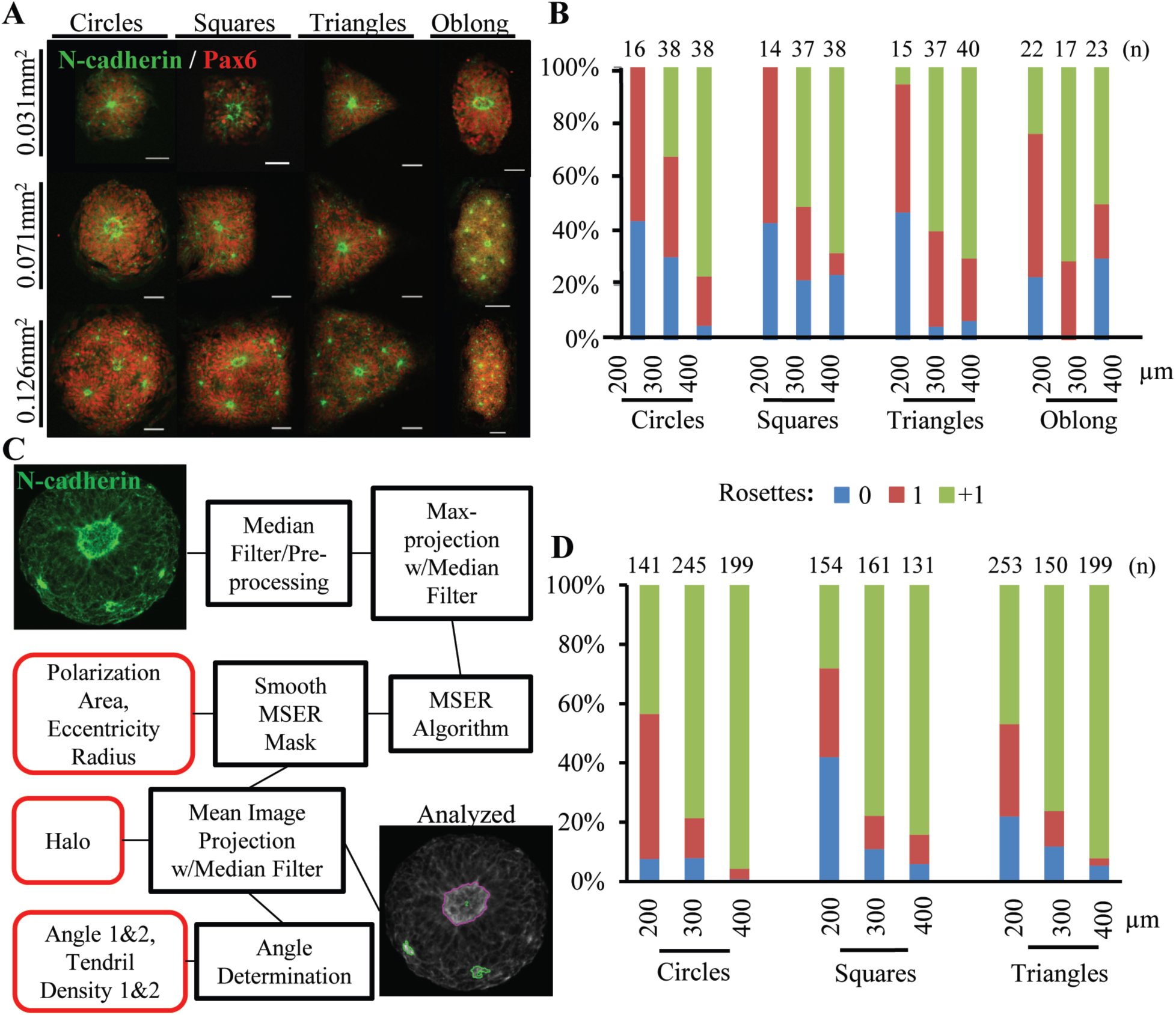
Effect of Tissue Morphology on Singular Rosette Emergence. (A)Representative images of micropatterned neuroepithelial tissues of circle, square, triangle, and oblong morphologies of varying size. Scale bars are 50μm. (B) Manual quantification of polarization foci and neural rosettes within micropatterned neuroepithelial tissues of varying morphology. Number of tissues analyzed per geometry is indicated above each bar. (C) Schematic of automated image analysis algorithm for quantification of polarization foci and neural rosettes. Purple outline indicates a rosette and green outlines indicate polarization foci as determined by the image analysis workflow. (D) Automated quantification of polarization foci and neural rosettes in micropatterned neuroepithelial tissues of varying morphology. Number of tissues analyzed per geometry is indicated above each bar.

To increase sample size, an automated workflow consisting of an image analysis algorithm and machine learning classifier was developed to quantitatively characterize and classify N-cadherin^+^ polarization foci and rosettes within micropatterned tissues (**Fig. 1C** and **Fig. S2**). The previous experiment was repeated using circular, triangle, and square micropatterns. Oblong micropatterns were omitted given their increased propensity to induce ‘+1 Rosette’ neuroepithelial tissues (**Fig. 2B**). Confocal image stacks (60X magnification) of Pax6/N-cadherin immunostained neuroepithelial tissues were collected for each morphology. Using only the N-cadherin immunostain channel, the analysis algorithm identified all areas of intense, spatially concentrated staining above a minimal area threshold, i.e. N-cadherin polarization foci, for each tissue and calculated eight descriptors for each area: polarization foci area, fitted ellipse major axis length and eccentricity, halo intensity ratio, tendril density 1 & 2, and tendril angle incoherence 1 & 2 (**Fig. S2** and Supplemental Methods). Collectively, these descriptors form a “descriptor vector” for each tissue.

Next, the MATLAB^®^ Machine Learning Toolbox was used to train a classifier based on descriptor vectors from 70, randomly selected, 300μm diameter (0.071mm^2^) circular tissues images (32). In addition to the algorithm analysis, these images were also examined independently by five NEC culture experts, who were asked to quantify the total number of N-cadherin^+^ polarization areas (i.e. foci) and determine which should also be classified as rosettes. Agreement between 4 out 5 experts was required to designate any polarization foci or rosette as a ‘ground truth’. Then, the human ground truth and the algorithm’s descriptor data for each identified N-cadherin polarization area in the 70 tissues (235 total identified areas) was used to learn a classifier using logic regression. Comparison of each human expert’s analyses versus the consensus ground truth revealed a human polarization foci and rosette identification error rate of 21.19^±^3.35% and 13.05^±^2.74%, respectively (**Table S1**). In comparison, the classification function exhibited lower error rates of 14.35% and 8.86%, respectively. To further test the classifier, a second round of human expert and algorithm analyses was conducted on an additional 35 neuroepithelial tissues selected randomly and distributed evenly across all morphologies. The human experts and automated image analysis algorithm identified 148 N-cadherin polarization areas total. The human versus classifier polarization and rosette error rates were 19.05^±^14.61% vs. 20.27% and 9.93^±^6.96% vs. 19.56%, respectively (**Table S2**). This suggest that the classifier generalizes well to new morphologies although the classifier’s rosette identification error rate increased potentially due to the variance in tissue morphologies. Still, the image analysis algorithm and classifier workflow was deemed acceptable for detecting trends in singular rosette emergence given the magnitude of differences observed in Figure 2B.

The automated workflow was used to analyze a total of 1633 images of neuroepithelial tissues with circular, triangle, and square morphologies across 0.031, 0.071, and 0.126mm^2^ surface area size scales. Compared to manual assessment, image analysis using the automated workflow estimated lower percentages of singular rosette emergence efficiency across all tissue morphologies (**Fig. 2D**). This was expected since the classifier’s rosette identification error rate is primarily due to false positives, i.e. erroneously classifying a polarization foci as rosette (**Table S2**). However, the trends in single rosette emergence efficiency were consistent with manual assessment in Figure 2B. The probability of biomimetic, singular rosette emergence increased as the neuroepithelial tissue’s surface area decreased. Interestingly, the automated analysis of increased sample size suggests that 200μm diameter (0.031mm^2^) circular versus square or triangle morphologies is more efficient at inducing singular rosette emergence, i.e. 48.9% vs. 29.9% vs. 31.2% respectively (**Fig. 2D**).

### Seeding at the hESC state increases singular rosette emergence with forebrain neuroepithelial tissues

Using the 200μm diameter (0.031 mm^2^) circular morphology, we re-examined whether the stage of neural induction at which cells are seeded onto micropatterned substrates affects the efficiency of singular rosette emergence. Our prior results indicated that rosette emergence was correlated with increasing cell density. Also, a significant increase in neuroepithelial cell density was noted between tissues formed using Pax6^−^ D1-2 cultures versus Pax6^+^ D3-5 cultures (**Fig. S1B**). Thus, we hypothesized that rosette emergence could be affected by the micropatterned cells’ proliferative capacity over 5 days of E6 culture. This was tested using the disparate cell phenotypes encountered during E6 neural induction, i.e. Pax6^−^ hESCs (D^−^1) and Pax6^+^ D4 NECs (**Fig. 3A**). Both cell types were seeded onto 200μm diameter (0.031mm^2^) circular micropattern arrays in biological duplicate, fixed, and immunostained after 5 days (**Fig. 3B**). In all tissues, the cells obtained a Pax6^+^/Otx2^+^ dorsal forebrain phenotype (23) with near uniformity. Manual analysis of over 100 neuroepithelial tissues in each experiment revealed a singular neural rosette emergence efficiency of 80.0±0.05% vs. 38.6±4.7% for neuroepithelial tissues formed by seeding hESCs vs. D4 NECs, respectively (**Fig. 3C**). The average cell density of neuroepithelial tissues formed using hESC vs. D4 NECs was ∼1.25 fold higher after 5 days of E6 culture or ∼ 30 more cells per tissue, i.e. 149 ± 23 vs. 119 ± 18 cells respectively (**Fig. 3D**). Although all tissues were multilayered, this change in cell density correlated with differences in neuroepithelial tissue morphology. Singular rosettes tissues derived from D^−^1 hESCs exhibited apico-basal polarity from a wider, central N-cadherin ring, while D4 NEC-derived singular rosette tissues had a tighter N-cadherin ring and appeared to have contracted slightly from their initial boundary (**Fig. 3B**, **E** and **F**). Hence, E6 culture of hESCs on 200μm diameter (0.031mm^2^) circular micropatterns induces highly reproducible, singular rosette emergence within forebrain neuroepithelial tissues in part due to their increased proliferative capacity.

**Fig. 3.**
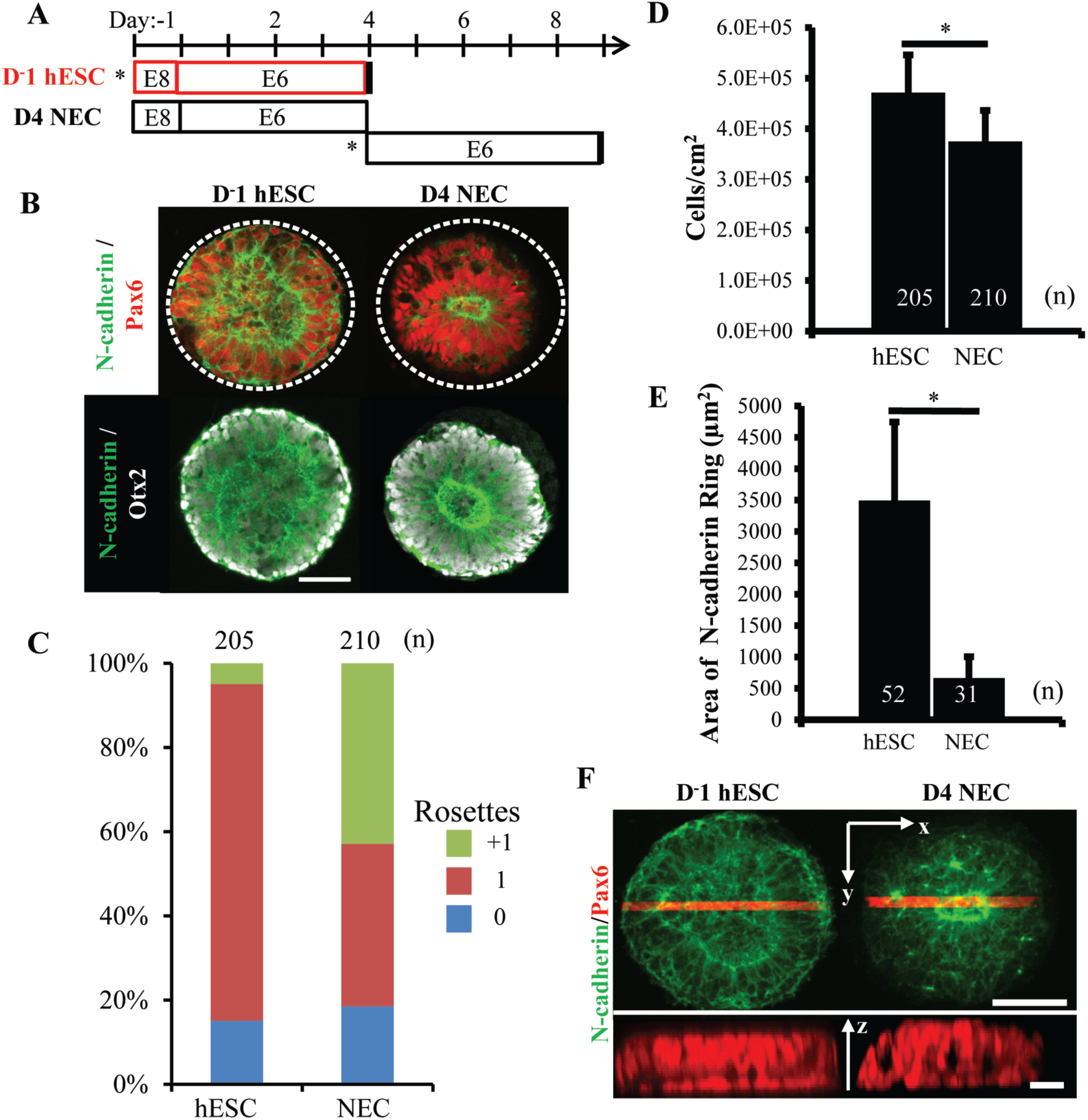
Comparison of Rosette Emergence in D^-^1 hESC vs. D4 NEC micropatterned tissues. (A) Schematic of micropatterned tissue derivation for D^-^1 hESCs and D4 NECs; sub-culture/seeding onto micropatterned substrates indicated by (*). (B) Representative immunostained images of forebrain, micropatterned neuropithelial tissues. (C) Manual quantification of polarization foci/neural rosettes per tissue with the number of tissues analyzed per condition indicated above each bar. (D) Average cell density and (E) area of polarized N-cadherin ring within micropatterned neuroepithelial tissues derived from D^-^1 hESCs and D4 NECs. Number of tissues analyzed per condition indicated on each bar, and error bars represent standard deviation. (*) indicates a significance of p<0.05 calculated using a One-way ANOVA. (F) Representative confocal Z-stack images of micropatterned neuroepithelial tissues (top) with profile view of red highlighted region (bottom). Scale bars are (B, F-top) 100μm and (F-bottom) 10μm.

### Controlled induction of singular neural rosettes within spinal tissues

The NEC polarized morphology is consistent throughout all regions of the primordial neural tube. However, their gene expression profile induced by morphogenetic patterning varies along both the neural tubes’ rostrocaudal (R/C) (33) and dorsoventral (D/V) (34, 35) axes. NECs derived using the E6 protocol patterns a default Pax6^+^/Otx2^+^ dorsal forebrain phenotype both in well plate and micropatterned tissue culture (**Fig. 3B**). We have also published a protocol for deriving NECs of discrete hindbrain through spinal cord regional phenotypes via deterministic patterning of their *HOX* expression profiles (36) (**Fig. 4A**). The protocol begins with high density seeding of hPSCs (D^-^1) followed by induction of a stable neuromesodermal progenitor (NMP) phenotype using sequential activation of FGF8b and Wnt signaling. Upon addition of the Wnt agonist (CHIR), *HOX* expression activates in a colinear and combinatorial manner over 7 days. At any time point within those 7 days, the NMPs can be differentiated into NECs by transitioning from a FGF8b/CHIR to a retinoic acid (RA) supplemented E6 media. Based on the time point at which this transition is made, the resulting NECs will express a distinct *HOX* profile indicative of phenotypical patterning to a discrete hindbrain through spinal cord R/C region. To enable highly reproducible derivation of biomimetic neural tissues from forebrain through spinal cord regions, we investigated how to integrate the *HOX* patterning protocol with the micropatterning methodology for inducing singular neural rosette emergence.

**Fig. 4.**
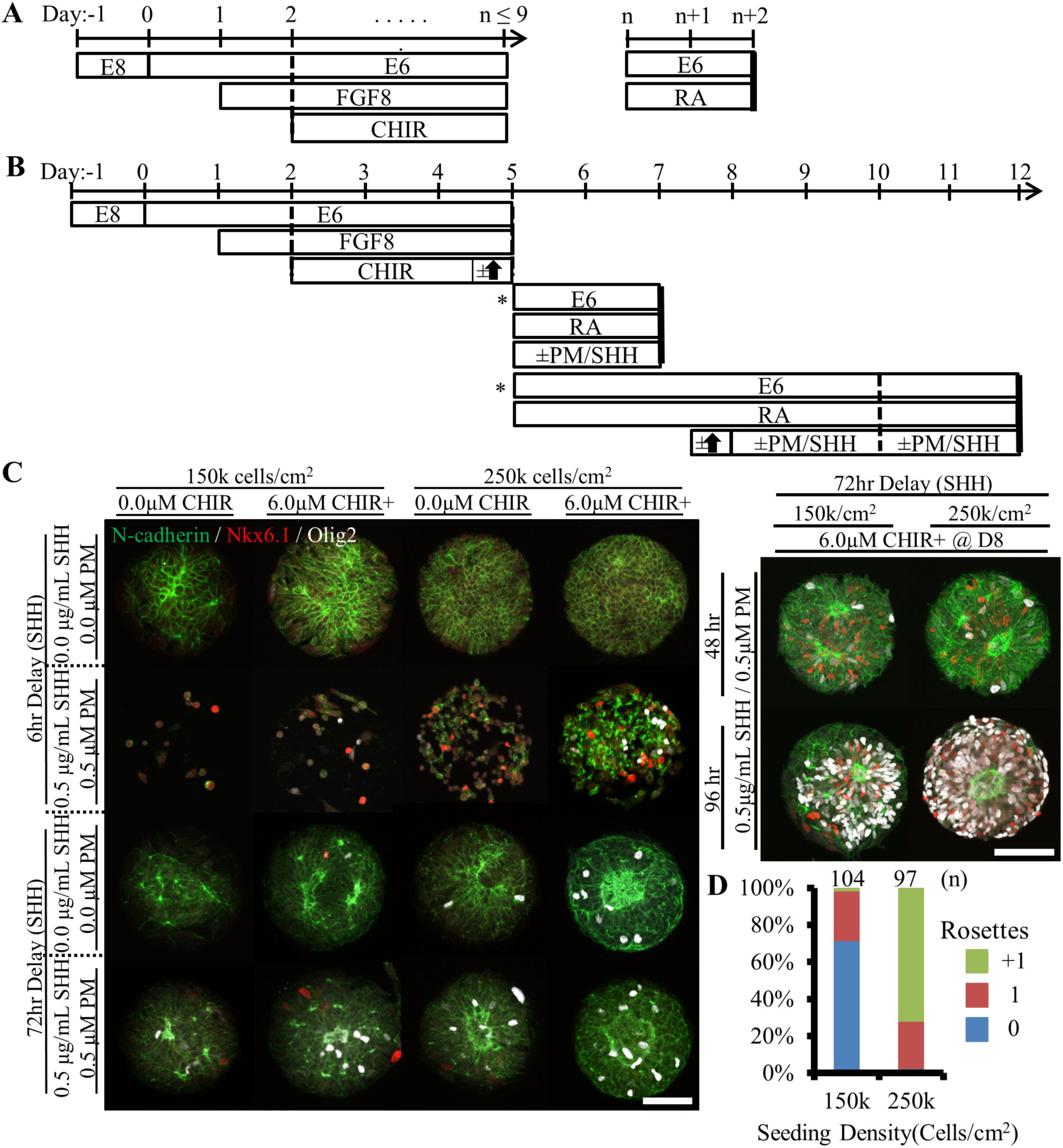
Derivation of Micropatterned Spinal Neuroepithelial Tissues. (A) Schematic for deterministic *HOX* patterning of hPSC-derived neuroepithelium in well plate culture. (B) Schematic for translation of well plate protocol to micropatterned arrays with subculture (*) of cervical *HOX* patterned NMPs to micropatterned substrates and subsequent differentiation to ventral spinal tissues. (C) Representative images of tissues derived when varying seeding density, timing of CHIR boost, and timing and duration of ventral patterning using Sonic Hedgehog (SHH/PM) signaling. Scale bars are 100μm. (D) Quantification of polarization foci/rosettes per tissue with the number of tissues analyzed per condition above each bar.

The mitogenic properties of FGF8 and Wnt prevent direct transfer of the *HOX* patterning protocol to D^-^1 hESC-seeded micropatterned substrates. After only a few days, the cells would proliferate to such a great extent that they would form spheroids that detached from the culture surface. Given our prior positive results with seeding Pax6^-^ cells (**Fig. 1H-I** and **Fig. 3**), we explored whether seeding Pax6^-^/Sox2^+^/T^+^ NMPs instead would still permit highly robust neural induction and singular rosette emergence. In this manner, the NMP culture could be caudalized to the desired R/C position using standard 6-well plate culture, and then seeded onto micropatterned substrates concurrent with transitioning to RA supplemented media to generate NECs (**Fig. 4B**). For these experiments, we derived spinal NECs using a 72-hour caudalization (i.e. CHIR exposure) period, which is known to pattern a lower brachial cervical/thoracic regional phenotype (36). Regional patterning was confirmed by qRT-PCR (**Fig. S3A**). Additionally, the ability to induce dorsoventral patterning with a transient spike in Wnt followed by sustained Sonic Hedgehog (SHH) signaling during neural induction was also tested. Both the transient Wnt (37) and sustained SHH (38) signaling are known regulators of ventral spinal patterning in vivo.

Using a combinatorial experimental design, the following variables were tested with spinal NMPs on 200μm diameter (0.031mm^2^) circular micropatterns: [1] NMP seeding density; [2] the application and timing of a transient Wnt (CHIR) boost; [3] timing of initiating ventral patterning, i.e. adding SHH and Purmorphamine (PM), a SHH agonist; [4] the duration of SHH ventralization (**Fig. 4B-C**). Rosette formation and ventralization were assessed by N-cadherin and Nkx6.1/Olig2, co-markers of spinal motor neuron progenitors, immunostaining respectively (23). Manual tissue analysis in each condition revealed that the optimal protocol regimen consisted of seeding spinal NMPs at 2.5×10^5^ cells/cm^2^ onto micropatterned substrates, culturing in RA supplemented E6 media for 72-hrs, and applying a 6-hr, 6.0μM Wnt boost immediately prior to 96-hrs of 0.5μg/mL SHH and 0.5μM PM exposure (**Fig. 4C**). Waiting 72-hrs after NMP seeding before applying the Wnt boost followed by SHH/PM exposure was the only tested regimen that consistently yielded both good cell viability and tissue formation as well as effective ventral patterning, i.e. 86±11% (n=8) Nkx6.1^+^/Olig2^+^ (**Fig. 4C and Fig. S3B-C**). Yet, only ∼25% (n=97) of neuroepithelial tissues on 200μm diameter (0.031mm^2^) circular micropatterns displayed singular rosette emergence (**Fig. 4D**).

### NEC biomechanical properties vary by regional phenotype and affect rosette emergence behavior

The inefficient induction of singular rosette emergence in spinal neuroepithelial tissues under micropatterning conditions that were highly efficient for forebrain NECs prompted a re-examination of the optimal circular micropattern size for each regional phenotype. Moreover, we investigate how altering the biomechanical properties of each regional neuroepithelial tissue, by partial inhibition of the Rho-associated protein kinase (ROCK), would affect which micropattern size induced optimal singular rosette emergence. D^-^1 hESCs and 72-hr caudalized NMPs were seeded at 100,000 and 250,000 cells/cm^2^, respectively, onto 150 (0.018mm^2^), 180 (0.025mm^2^), 200 (0.031mm^2^), 250 (0.049mm^2^), and 400μm (0.126mm^2^) diameter circular micropattern arrays to generate forebrain and spinal neuroepithelial tissues (**Fig. 5A-B and D-E**). After 5 days of culture in the absence or presence of 10μM ROCK inhibitor, manual analysis of rosette emergence revealed stark differences in the emergent behavior of forebrain vs. spinal neuroepithelial tissues. Similar to our previous results, the efficiency of singular neural rosette emergence for forebrain tissues peaked at 85% on 250μm (0.049mm^2^) diameter circular micropatterns (**Fig. 5C**). However, in the presence vs. absence of ROCK inhibitor, the efficiency curve shifted leftward inducing singular rosette emergence in 75% vs. 0% of 150μm(0.018mm^2^), 89% vs. 50% of 180μm (0.025mm^2^), 87% vs. 58% of 200μm (0.031mm^2^), and 11% vs. 85% of 250μm (0.049mm^2^) diameter forebrain neuroepithelial tissues, respectively. In comparison, singular neural rosette emergence within spinal neuroepithelial tissues peaked at 73.5% on 150μm (0.018mm^2^) diameter micropatterns, and the presence of ROCK inhibitor completely abolished their ability form neural rosettes (**Fig. 5F**). Of note, NEC density within forebrain vs. spinal tissues of a given micropattern size and in the presence vs. absence of ROCK inhibitor did not vary significantly (**Fig. S4**).

**Fig. 5.**
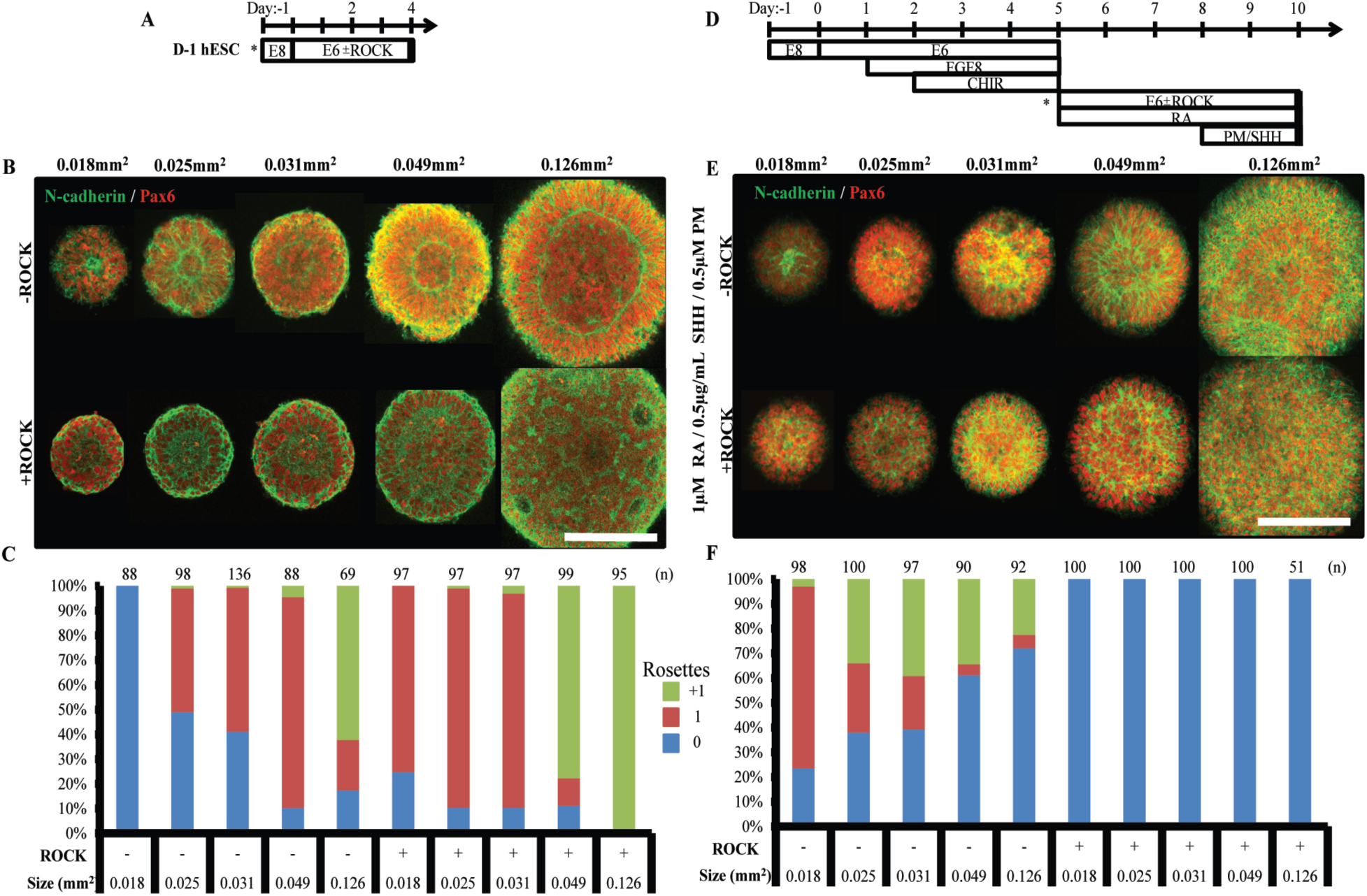
Singular Neural Rosette Emergence within Forebrain and Spinal Neuroepithelial Tissues. (A) Schematic for derivation of micropatterned forebrain neuroepithelial tissues; sub-culture/seeding onto micropatterned substrates indicated by (*). (B) Representative images of neural rosette emergence in micropatterned forebrain neuroepithelial tissues of various areas with and without ROCK inhibitor. (C) Quantification of polarization foci/rosettes per forebrain tissue with the number of tissues analyzed per condition above each bar. (D) Schematic for derivation of micropatterned spinal neuroepithelial tissues; sub-culture/seeding onto micropatterned substrates indicated by (*). (E) Representative images of neural rosette emergence in micropatterned spinal cord neuroepithelial tissues of various areas with and without ROCK inhibitor. (F) Quantification of polarization foci/neural rosettes per spinal tissue with the number of tissue analyzed per condition above each bar. Scale bars are (B, F) 200μm.

Taking into account our previous experiment (**Fig. 3**), the optimal morphology for single neural rosette emergence in forebrain and spinal neuroepithelial tissues is 200 - 250μm (0.031 - 0.049mm^2^) and 150μm (0.018mm^2^) diameter circular micropatterns respectively. The physical size of emergent neural rosettes is not conserved across tissues of increasing area (**Fig. 5B and E**). Hence, the micropatterned tissue morphology does not instruct rosette morphogenesis, but restricts tissue area to statistically favor emergence of a single rosette in accordance with innate NEC properties. Since ROCK inhibition down-regulates actin polymerization and acto-myosin contraction (39), the results in Figure 5 imply that decreasing the contractility of forebrain NECs reduces the length scale over which the cells can effectively polarize to evoke a rosette structure, i.e. 250μm vs. 150/180/200μm in 0 vs. 10μM Rock inhibitor. This further suggests that forebrain vs. spinal NECs can generate more contractile force since singular rosette emergence peaks in larger forebrain, i.e. 250μm (0.049mm^2^) diameter, versus smaller spinal, i.e. 150μm (0.018mm^2^) diameter, circular tissues. The ability of 10μM Rock inhibitor to only partially diminish vs. fully eliminate the rosette formation capacity of forebrain vs. spinal NECs also supports this ranking of contractility. These results provide the first evidence that NEC biomechanical properties vary based their regional patterning. They also demonstrate the necessity of customizing the micropatterned neuroepithelial tissue morphology to efficiently induce singular rosette emergence based on the cell’s regional phenotype.

### Singular neural rosette cytoarchitecture is maintained during radial tissue outgrowth

The importance of maintaining a singular neuroepithelium in early CNS development is evident given severe congenital disorders associated with breakdown (40, 41) or duplication (12, 14) of the neuroepithelial tube. Accordingly, we investigated whether the singular rosette structure induced within micropatterned neuroepithelial tissues would persist upon subsequent tissue growth and morphogenesis. Using our previously developed ‘clickable’ polyethylene glycol (PEG) brush chemistry (26, 28), we synthesized array substrates that could be actuated to release the micropatterned tissues from their confinement and permit radial tissue outgrowth. In our prior publication, tissue outgrowth was enabled by in situ conjugation of-RGD peptide bioconjugates to the microarray’s PEG brushes. However, this caused aberrant migration of single cells away from the neuroepithelial tissues instead of a more coordinate, biomimetic radial tissue expansion (26). Therefore, we explored whether a clickable bioconjugate containing a heparin-binding peptide (-CGTYRSRKY) derived from Fibroblast growth factor-2 would mediate a more gradual tissue expansion. This was posited based on the fact that the heparin-binding peptide (HBP) is unable to directly promote cell migration via integrin binding but could immobilize heparin sulfate proteoglycan-bound ECM proteins (e.g. Laminin) secreted at the neuroepithelial tissues’ basal aspect upon extended culture (42) (**Fig. S5**).

To evaluate feasibility of our proposed substrate design, micropatterned arrays with reactive, 200μm (0.031mm^2^) diameter, circular PEG brushes were fabricated and functionalized via an overnight incubation in cell culture media containing 0 to 20**μ**M FITC-HBP conjugated to a Malemide-PEG4-DBCO linker (**Fig. S6 and S7A-B**). Using confocal microscopy, a direct correlation between fluorescence from immobilized FITC-HBP and the incubation media concentration was observed with culture media containing 20**μ**M FITC-HBP-DBCO bioconjugate providing maximal surface density. Then, we tested whether HBP-presenting PEG brushes could sequester ECM proteins. Substrates modified with cell culture media containing 0, 5, 10, and 20**μ**M HBP-DBCO bioconjugate were subsequently incubated overnight in media containing 10**μ**M NHS-rhodamine tagged Laminin, which was isolated from a mouse sarcoma and likely cobound with heparin sulfate proteoglycans. Imaging revealed a consistent increase in rhodamine fluorescence directly correlated with increasing surface densities of HBP and saturating on substrates modified with culture media containing 10**μ**M HBP-DBCO bioconjugate (**Fig. S7C-D**). Thus, PEG-brushes modified in situ with 10**μ**M HBP-DBCO bioconjugate are capable of immobilizing Laminin proteins, and potentially a diverse cadre of basement membrane ECM proteins that bind heparin sulfate proteoglycans (43).

Next, neuroepithelial tissue outgrowth was tested on both inert and reactive PEG-grafted substrates micropatterned to display arrays of 200μm (0.031mm^2^) diameter circular culture regions with 400μm center-to-center spacing that were subsequently coated with Matrigel. Pax6^-^ D^-^1 hESCs were seeded onto the substrates at 100,000 cells/cm^2^ and maintained in E6 culture for 3 days, at which point the media was supplemented with 10.0μM HBP-DBCO bioconjugate for 24 hours on both inert and reactive microarray substrates. After 3 additional days of culture, the tissues were fixed and stained for N-cadherin and Laminin (**Fig. 6A**). Inert control PEG brushes predictably maintained the restricted singular rosette tissue morphology for the culture duration (**Fig. 6B**). In contrast, on reactive substrates modified in situ with HBP-DBCO bioconjugates, the tissues expanded as a coherent, multilayered structures maintaining a singular neuroepithelium and eventually intersecting with neighboring tissues. Also, they displayed basal deposition of laminin resembling the developing neural tube’s basal lamina (**Fig. 6C-D**). To prevent inhibition of tissue outgrowth, the experiment was repeated on reactive micropatterned arrays with 200μm (0.031mm^2^) diameter circular regions with larger 800μm center-to-center spacing. The in situ modification occurred on Day 4 of E6 culture, and microarrays were fixed and stained after an additional 7 days of culture (**Fig. 6E**). These tissues also maintained a singular Pax6^+^ neuroepithelium throughout the culture period that morphed into a 3-D hemispherical structure with a central cavity and Tuj1^+^ neuronal cells residing at the tissues periphery (**Fig.S7 and 6F-G**). These results indicate that controlled induction of singular rosette emergence upon organotypic neural tissue formation will persist throughout subsequent stages of morphogenesis, thereby enabling formation of a biomimetic nascent CNS tissue cytoarchitecture.

**Fig. 6.**
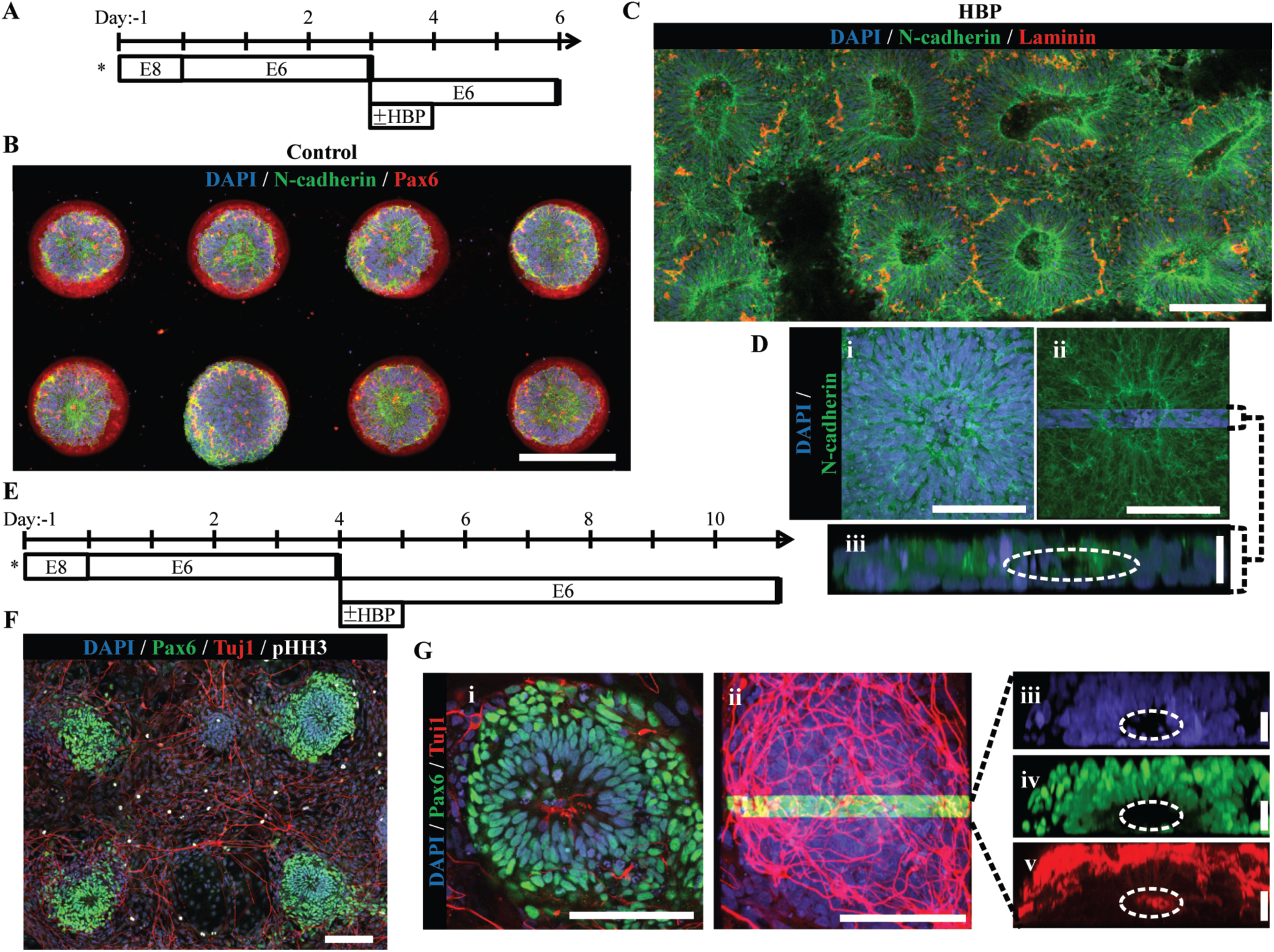
Micropatterned Neuroepithelial Tissue Outgrowth on Clickable, HBP-presenting Substrates. (A) Schematic for derivation and HBP-mediated expansion of forebrain neuroepithelial tissues on chemically-modified micropattern arrays of 200μm diameter circles with 400μm center-to-center spacing. (B) Image of singular neural rosette emergence within D6 neuroepithelial tissues on inert PEG-MA substrates not functionalized by HBP-DBCO bioconjugates. (C) Image of singular neural rosette expansion within D6 neuroepithelial tissues on reactive PEG-MA substrates functionalized with HBP-DBCO bioconjugates. The tissues show radial expansion and deposition of a Laminin^+^ basement membrane. (D) (i) Magnified image of expanded neuroepithelial tissue with (ii) highlighted 3D cross-section and (iii) profile view; white dotted line indicates hollow polarization cavity. (E) Schematic for derivation and HBP-mediated expansion of forebrain neural tissues on chemically-modified micropattern arrays of 200μm diameter circles with 800μm center-to-center spacing. (F) Image of singular neural rosette tissue outgrowth and neuronal differentiation on HBP-functionalized substrates at day 11. (G) (i) Magnified image of the neuroepithelial center of expanded neural tissues with (ii) the highlighted 3D cross-section and profile view of (iii) nuclei, (iv) Pax6^+^ core, and (v) primarily peripheral Tuj1^+^ neuronal processes; white dotted lines indicate hollow polarization cavity. Scale bars are (B, C) 200μm, (D(i,ii), G(i,ii)) 100μm, and G(iii,iv,v)) 20μm.

## Discussion

Neural organoid technology has enabled unprecedented in vitro recapitulation of biomimetic, human CNS tissue microenvironments. This is currently being used to develop novel personalized neurological disease modeling platforms (4, 44, 45). However, while impressive in their extent of ex vivo morphogenesis, the spontaneous and cell-intrinsic emergent properties that enable organoid formation also currently limits their wide-scale implementation due to the lack of reproducibility in their macroscale cytoarchitecture and cellular composition. In essence, we now have a glimpse of the neural organoid’s potential, but do not understand the processes that govern their morphogenesis well enough to precisely engineer their formation (46). Here, we have melded organoid technology with a biomaterial culture platform to elucidate biophysical parameters that control the genesis of human neural organoid morphogenesis, i.e. polarized neural epithelium formation.

Over the past decade, scientists have generated human neural organoids of the cerebral cortex (2), retina (3), forebrain (4, 17), midbrain (47), and cerebellum (48) with both neuronal and glial cellular constituents (49). However, a consistent limitation in creating reproducible and biomimetic neural organoid anatomy at the macroscale is the random formation of multiple polarized neuroepithelial regions, a.k.a. rosettes, which can each act as independent morphogenesis centers. Using micropatterned culture substrates, we have demonstrated that enforcing a 200-250μm forebrain and 150μm spinal, circular, neuroepithelial tissue morphology optimally induces formation of a singular rosette cytoarchitecture with 80-85% and 73.5% efficiency respectively (**Fig. 3 and 5**). Our results indicate that these biophysical parameters for reproducible induction of a biomimetic, neural tube-like cytoarchitecture are dependent on local cell density, acquisition of Pax6 expression, and cell non-autonomous biomechanical properties such as contractility. Furthermore, we have shown that the singular neural rosette cytoarchitecture can be maintained upon subsequent tissue morphogenesis, indicating that initial constraint of tissue morphology in accordance with these identified length scales could be a generalizable approach to engineering human neural organoids with standardized nascent CNS tissue cytoarchitecture. However, we acknowledge that our micropatterned tissue do morph from 2-D monolayers into 3-D hemispherical tissues (**Fig. 6**), and therefore, our identified optimal tissue morphology parameters may not be precisely conserved for organoids initially formed as 3-D cell aggregates.

While organoids can be used to generate novel humanoid disease models, their ability to recapitulate developmental morphogenesis ex vivo and within a human genomic background also provides unique opportunities to investigate mechanisms that orchestrate early human development (21, 22). In efforts to develop a unified approach for inducing singular neural rosette emergence within neuroepithelium patterned to any CNS rostrocaudal region, we discovered that the biomechanical properties of Pax6^+^/N-cadherin^+^ NECs of the dorsal forebrain differed from those of the ventral spinal cord. Forebrain NECs appear to be able to form singular neural rosette within tissue of larger circular dimensions than spinal NECs because of their increased contractility (**Fig. 5**). This would seem to correlate with the significantly larger dimensions of forebrain ventricles versus the spinal cord’s central canal, and could represent a novel axis between the biochemical morphogens and biomechanical properties that govern CNS morphogenesis. While our conclusions are based on observations from correlated perturbations of ROCK activity and not direct contractility measurements, our experimental paradigm does provide a unique platform for further definitive analysis of the interplay between biochemical and biomechanical factors that govern this facet of human CNS morphogenesis. To our knowledge, there is no other experimentally tractable means by which such phenomena can be investigated within a human genomic background.

The ability to reproducibly engineer the nascent neuroepithelial cytoarchitecture of neural organoids in a high-throughput arrayed format could have significant implications for future advanced biomanufacture of the neural organoid platform. The 3-D nature of organoids grown in suspension culture provides constituent cells with a biomimetic tissue context. However, suspension organoids grow to millimeters in diameter limiting real-time analysis of internal morphogenesis processes using traditional imaging modalities. While our microarrayed neural tissues initiate as a 2-D monolayer, they quickly become multilayered and definitively 3-D, i.e. >4 cell layers thick, within 4 days of culture and continue to increase in thickness upon extended culture (**Fig. 3** and **6**). Thus, the biophysical (50) and electrophysiological (51) benefit of 3-D organoids may still apply within our engineered neural tube slides. Our arrayed approach enables constant monitoring throughout their morphogenesis process using confocal microscopy. Furthermore, the stratified microscale cytoarchitecture observed in regions of cortical, retinal, cerebellar, and cerebral organoids may also be achievable upon further modification of our culture substrates to provide continual but expanding confinement as the arrayed tissues outgrow radially (26, 52). If achievable, then the substrate parameters elucidated in this study could serve as a basis for generating an engineered well-plate platform to which scientists could simply add their patient-specific hPSCs or Pax^-^ hPSC derivatives (**Fig. 3** and **4**), follow an in situ modification protocol (**Fig. 6**), and allow the engineered culture substrate to instruct reproducible growth of high density arrays of neural organoid tissues. This would facilitate broad dissemination of the neural organoid platform as well as real-time microscopic investigation of the organoid’s morphogenesis and terminal structure.

## Methods

### Micropatterned Array Substrate Fabrication

Micropatterned array cell culture substrates were fabricated using a combination of previously published methods (26, 28). Polydimethylsiloxane (PDMS) stamps with arrays of post and micro-well features were generated as relief molds of silicon wafers. The wafers were designed in AutoCAD and purchased from FlowJEM (www.flowjem.com). PDMS stamps were coated with ω-mercaptoundecyl bromoisobutyrate (2mM in 100% ethanol), dried under inert gas, and then brought in conformal contact with glass coverslips coated with 180nm Au atop 30nm Ti. Micropatterned slides were then incubated in 100% ethanol for 10 minutes, prior to being dried under nitrogen and transferred to a Schlenk flask under vacuum. A solution of either poly(ethylene glycol) methyl ether methacrylate (PEG-MEMA) or poly(ethylene glycol) methacrylate (PEG-MA) macromonomer (Sigma Aldrich) with water, methanol (Thermo Fisher), copper(II) bromide (Sigma Aldrich), and 2’,2-bipyridine (Sigma Aldrich) was degassed and transferred to the reaction flask. Surface-initiated atom-transfer radical-polymerization (SI-ATRP) of PEG polymers was initiated by injection of L-ascorbic acid (Sigma Aldrich) in deionized water into the reaction flask. ATRP was allowed to continue for 16 hours at room temperature to generate micropatterned PEG brushes. Polymerization was terminated via addition of air and followed by rinsing with ethanol and water before drying under inert gas. In a sterile hood, the substrates were rinsed five times with sterile PBS (Thermo Fisher) and transferred to individual wells of a 12-well tissue-culture polystyrene (TCPS) plate where they were rendered cell-adhesive through adsorption of 0.083mg/mL Matrigel (WiCell) in DMEM/F-12 (Thermo Fisher) via overnight incubation at 37°C.

### Generation of Micropatterned Forebrain Neuroepithelial Tissues

WA09 or HUES3 Hb9::GFP hESCs were maintained in Essential 8 medium (E8) on Matrigel-coated TCPS plates and routinely passaged with Versene (Thermo Fisher). NEC derivation from hPSCs was performed in accordance with the E6 protocol (23). To generate NEC-derived micropatterned tissues at various stages of neural derivation, cells were first rinsed with PBS, dissociated with Accutase (Thermo Fisher) for 5 minutes at 37°C, and collected via centrifugation at 1000rpm for 5 minutes. Singularized NECs were re-suspended in E6 media with 10μM ROCK inhibitor (Y27632; R&D Systems) and seeded onto micropatterned substrates at 75,000 cells/cm^2^ in 2mL of media per well. The following day the media was replaced with 2mL of E6 media and 50% media changes were performed daily thereafter. Alternatively, to generate hPSC-derived micropatterned tissues, hPSC cultures at ∼85% confluency were rinsed with PBS, dissociated with Accutase for 5 minutes at 37°C and collected via centrifugation at 1000rpm for 5 minutes. Singularized hPSCs were then suspended in E8 media with 10μM ROCK inhibitor and seeded onto micropatterned substrates at 100,000 cells/cm^2^ in 2mL of media per well. The following day the media was replaced with 2mL of E6 media and 50% media changes were performed daily thereafter.

### Generation of Micropatterned Spinal Cord Neuroepithelial Tissues

Caudalization of hPSC-derived cultures to generate micropatterend neuroepithelial tissues of the cervical/thoracic spinal cord was performed using a modified version of the deterministic *HOX* patterning protocol (36). Human PSC’s were dissociated with Accutase and seeded onto matrigel-coated, 6-well TCPS plates at 150,000 cells/cm^2^ in E8 media with 10μM ROCK inhibitor. The following day the media was switched to E6 media for 24hrs, after which the media was supplemented with 200 ng/mL FGF8b (R&D Systems) to initiate differentiation into a NMP phenotype. Following 24hrs in E6^+^FGF8b media, the cells were dissociated with Accutase for 1:45 minutes and seeded onto TCPS plates at a 1:1.5 well ratio in E6 media with 200ng/mL FGF8b, 3μM CHIR (R&D Systems), and 10μM ROCK inhibitor. The cells were allowed to remain in E6^+^FGF8b^+^CHIR^+^ROCK inhibitor for 48hrs before the media was switched to E6^+^FGF8b^+^CHIR. After 24hrs, the NMPs were dissociated with Accutase for 5 minutes and seeded onto micropatterned substrates at 150,000 cells/cm^2^ or 250,000 cells/cm^2^ in E6 media with 10μM ROCK inhibitor. Six hours later, the media was supplemented with 1μM ROCK Retinoic Acid (RA, R&D Systems) and 10μM ROCK inhibitor to initiate transition to a neuroepithelial cell type. The next day the media was switched to E6 media with 1μM RA and 50% media changes were performed every 24 hrs for the next 2 days. Then, the media was supplemented with 1μM RA, 2μg/mL Sonic hedgehog, and 2μM Purmorphomine (Shh, PM; R&D Systems) to induce ventralization. To further enhance ventralization, a 6hr boost in Wnt signaling via media supplementation with 6μM CHIR was applied either immediately before subculture or 72hrs after cell seeding onto micropatterned substrates but prior to RA+SHH+PM exposure. The neuroepithelial cells were maintained in E6+RA+SHH+PM with 50% media changes daily.

### In Situ Modification of Micropatterned Neural Tissues Arrays

Micropatterned substrates amenable to in situ click modification and the clickable bioconjugates were synthesized as previously described elsewhere (26). Detailed methods are provided in *Supporting Methods.* After seeding hPSCs to generate micropatterned neuroepithelial tissue arrays as previously described, E6 media supplemented with 10μM HBP-DBCO bioconjugate was added on day 3 or 4. After 24hrs, the media was replaced with 2mL E6 media and 50% media changes were performed daily thereafter.

### Quantification of Cell Density

Quantification of cell density in micropatterned neuroepithelial tissues was performed using image segmentation packages in CellProfiler (BROAD Institute) to identify and count DAPI stained nuclei in 3D image stacks. Image segmentation was also used to identify and quantify the number of nuclei stained positively for transcription factors Pax6, Otx2, Nkx6.1, and Olig2. The number of nuclei expressing the transcription factor of interest was compared to the total DAPI nuclei present to estimate percentages of regionally patterned NECs.

### Immunocytochemistry and Microscopy

Micropatterned tissues were rinsed with PBS and fixed with 4% paraformaldehyde (PFA, Sigma Aldrich) in PBS for 10 minutes at room temperature. Following three rinses with PBS, the micropatterned substrates were blocked and permeabilized in PBS with 5% Donkey Serum and 0.3% Trition X-100 (TX100; Thermo Fisher) (PBS-DT) for 1 hour at room temperature. Next, the tissues were incubated with primary antibodies diluted in PBS-DT for 2 hrs at room temperature or overnight at 4°C. Then, the substrates were rinsed in PBD-DT three times for 15 minutes each. Secondary antibodies were diluted in PBS-DT and exposed to the tissues for 1 hour at room temperature. A list of primary antibodies and dilutions can be found in **Table S3**. Tissue nuclei were subsequently stained with 300nM 4’,6-diamidino-2-pheny-lindoldihydrochloride (DAPI, Sigma Aldrich) for 30 minutes followed by three 15 minute PBS washes. The micropatterned substrates were mounted on glass coverslips using Prolong Gold Antifade Reagent (Thermo Fisher). Brightfield images of micropatterned tissues were obtained using a Nikon TS100 microscope with a ME600L camera. Fluorescence images were obtained using a Nikon A1R confocal microscope.

## Author Contributions

G.T.K. and R.S.A conceived the study. G.T.K. and B.F.L. performed all substrate synthesis. G.T.K. performed all cell-culture experiments. G.T.K., R.S.A., R.L.W., and W.S. designed the parameters for automated neural tissue analysis and R.L.W. and W.S. programmed the algorithm. L.M.T.A. developed the machine learning classifier. All authors contributed to manuscript preparation. The authors declare no conflicts of interest.

## Acknowledgements

This publication was developed, in part, using the Assistant Agreement No. 83573701 awarded by the U.S. Environmental Protection Agency (EPA) to R.S.A. and has not been formally reviewed by the EPA. The views expressed in this document are solely those of R.S.A. and colleagues and do not necessarily reflect those of the Agency, and the EPA does not endorse any products or commercial services mentioned in this publication. The publication was also developed with support from NIH grants R21NS082618 and R33NS082618, an Innovation in Regulatory Science Award from the Burroughs Wellcome Fund (R.S.A.), and seed project funding from the Wisconsin Institute for Discovery.

## Abbreviations

hPSC: human pluripotent stem cell
hESC: human embryonic stem cell
CNS: central nervous system
NEC: neuroepithelial cell
NMP: neuromesodermal progenitor
PEG: poly(ethylene glycol)
RA: retinoic acid
PM: purmorphamie
SHH: sonic hedgehog
FITC-HBP: fluorescein isothiocyanate-tagged heparin binding peptide
PEG4-DBCO: poly (ethylene glycol)_4_-dibenzocyclooctyne.
TCPS: tissue culture polystyrene

